# Online characterization of surrogate metrics for metabolic phenotype in human induced pluripotent stem cell bioprocessing

**DOI:** 10.64898/2026.05.08.723750

**Authors:** James Colter, Michael Scott Kallos, Kartikeya Murari

## Abstract

Human induced pluripotent stem cells (hiPSCs) are the most accessible source material for derivation of stem-cell-based therapies at scale. However, a disconnect exists between quality characteristics of phenotype in the pluripotent state, and downstream metrics for efficacy and safety. Bridging this gap is a major challenge. Given hiPSC plasticity, environmental conditioning plays a crucial role in guiding phenotype. This work presents a parallelizable scale-down approach, acquiring real-time data to inform hiPSC phenotype throughout biomanufacturing. We developed an optoelectronic instrumentation suite capable of measuring pH, dissolved oxygen, and cell density as important surrogates for phenotype in a scale-down expansion bioprocess. We were successful in obtaining continuous, integrated parametric data throughout cultivation and estimating metabolic characteristics of hiPSC phenotype. This system functions as a proof-of-concept tool for development of predictive models and monitoring strategies around the elucidation of phenotypic dynamics within hiPSC biomanufacturing. We have demonstrated a feasible open-source multivariate continuous monitoring approach at research scale that combines common process parameters with a scattering measurement against aggregate density. The combination of these parameters enables surrogate measurement of a metric for metabolic phenotype. This contribution emphasizes monitoring how the bioprocess influences variables important in the context of cell state, in broader pursuit of better understanding the link to downstream functionality and global optima in hiPSC biomanufacturing for regenerative medicine.

## Introduction

Human induced pluripotent stem cells (hiPSCs) are an optimal cell type for the derivation of therapeutics given their capacity for self-renewal and differentiation into any adult cell type within the primordial germ layers [1,2]. As these cells are reverted to pluripotency from a somatic or multipotent state, they can effectively be generated anywhere along the developmental timeline or from adult tissue. This provides unique opportunities for their utility in both allogeneic and autologous forms of therapy, drug development, and disease modeling [3–5]. However, iPSC-derived therapeutics form an extremely small cohort of clinical trials being carried out in the United States, Europe, and Japan, and transparency concerning biological characteristics and associated clinical outcomes is poor [6–9]. Aside from having a complex pipeline spanning from induction of pluripotency to differentiation to target phenotypes, clinical efficacy remains an obstacle to overcome.

There is a large body of evidence in literature to suggest that despite showing similar identities, iPSC-derived cell types do not function in entirely predictable ways or with the same level of competency compared to embryonic stem cells (ESCs). Not only do pluripotent cells have a very high degree of epigenetic plasticity, but somatic patterns of regulation continue to exist post-induction, contributing to the myriad factors regulating function and phenotype [10–15]. These complexities are exacerbated by donor-to-donor variability [16, 17]. Further, modulation of phenotype engages a complex network of activities, and our groups have argued previously that conventional methods for characterization of phenotype do not effectively encapsulate interactions within this network [8].

While omics and process datasets are generated to ensure quality control metrics are met industrially, data are underutilized and a disconnect exists between the environmental conditioning of the bioprocess pipeline and phenotypic outcomes in the resulting population [18]. Further, while systems exist to monitor process parameters throughout the process, these are generally designed for either large-scale commercial use or integrated culture platforms, making them cost-prohibitive for widespread generation of data in research and development environments encompassing a range of bioprocess configurations [19]. Given the significance of the properties of the culture vessel in determining culture dynamics and scalability, having phenotype metrics that are easily translatable across processes and researchers is critical in the context of global optimization of the bioprocess pipeline. The existing gaps have driven us to find ways to model interactions within the cells and across the population, in a modality that is well integrated with a scale-down process environment. A visualization of these high-level interactions is shown (Fig 1a).

**Figure 1.**
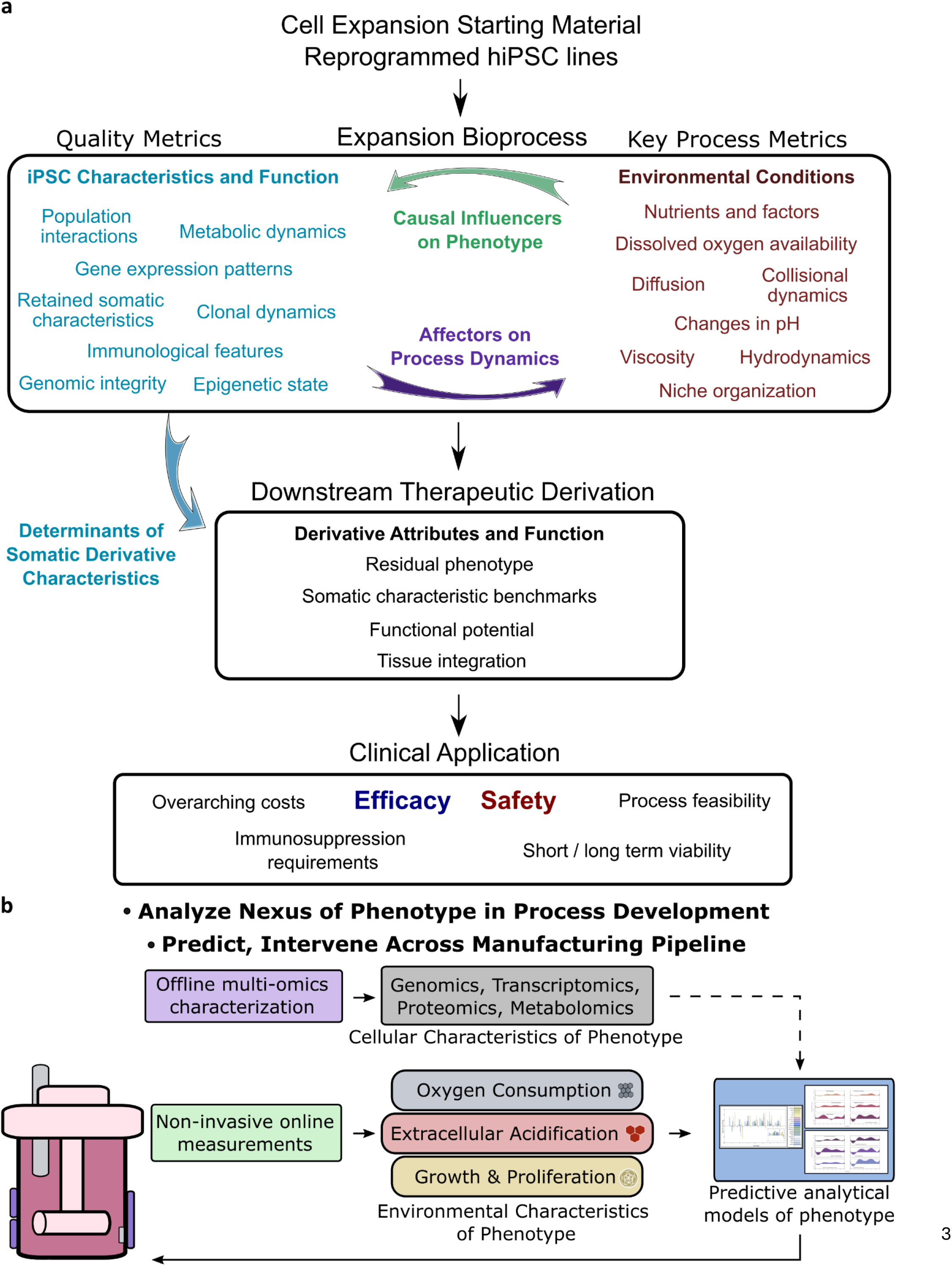
(a) Consolidation of key considerations for modulation of hiPSC phenotype within the process environment, and downstream implications. Feedback loops between iPSC characteristics and function, environmental conditions, and derivative attributes and function are shown to highlight the importance of complex artificial niche considerations for therapeutic derivation. (b) Visualization of how the optoelectronic system presented herein may be translated to integrated approaches to model and predict phenotype dynamics in iPSC bioprocessing. By combining surrogate metrics with robust offline datasets, predictive models for global optima concerning hiPSC phenotype may be developed.

Therefore, we considered it pertinent to implement an open-source modular acquisition system to acquire multiple critical process parameters simultaneously at a laboratory-scale that can be effectively combined with offline omics datasets to generate models for phenotypic interactions within the bioprocess. Several challenges exist to fulfill these requirements within the constraints of the bioprocess being used, such as parameters measured, minimal footprint to enable use with scale-down process systems, parallelizability across biological replicates, minimization of disruption of the bioprocess, low heat-generation, sterility, humidity-protection, resistance to environmental perturbations, long-term data reliability, and scalability across research systems. In this study we aimed to demonstrate continuous online optoelectronic measurement of pH, cell density, and dissolved oxygen concentration in parallel in real-time over the course of a scale-down hiPSC bioprocess passage. This approach and its proposed utility in the field is shown at a high level of abstraction (Fig 1b). These metrics were chosen as surrogate markers of metabolic phenotype, and for their proposed utility in informing cell state when combined with more advanced characterization methods [20, 21]. Oxygen was chosen as the modulatory parameter given growing evidence in literature for its importance in cell state transition dynamics, both in natural embryonic development and in vitro [22, 23].

We developed a modular system of USB-powered printed-circuit-boards (PCBs), comprised of a controller, sensor boards, and a serial-peripheral interface (SPI). Onboard reference voltage and temperature measurements were included for signal correction and protocol-gating – capturing disruptions to acquisition that were confirmed as the result of culture-protocol-associated handling and correcting relative measurements as a result – during data analysis. Optical interactions were chosen to maintain a minimally invasive, easily parallelizable approach to culture monitoring. We report this approach to monitoring surrogate measures of metabolic phenotype, evaluating the capabilities of the optoelectronic system using well-established principles for calculation of pH by application of Beer-Lambert law, cell density by modeling optical interactions with cell aggregates, and dissolved-oxygen fluorescence quenching described by Stern-Volmer relation (Fig 2). Further, we report application of the system to study the feasibility of measuring surrogate metabolic phenotype for scale-down hiPSC bioprocesses under differential oxygenation.

**Figure 2.**
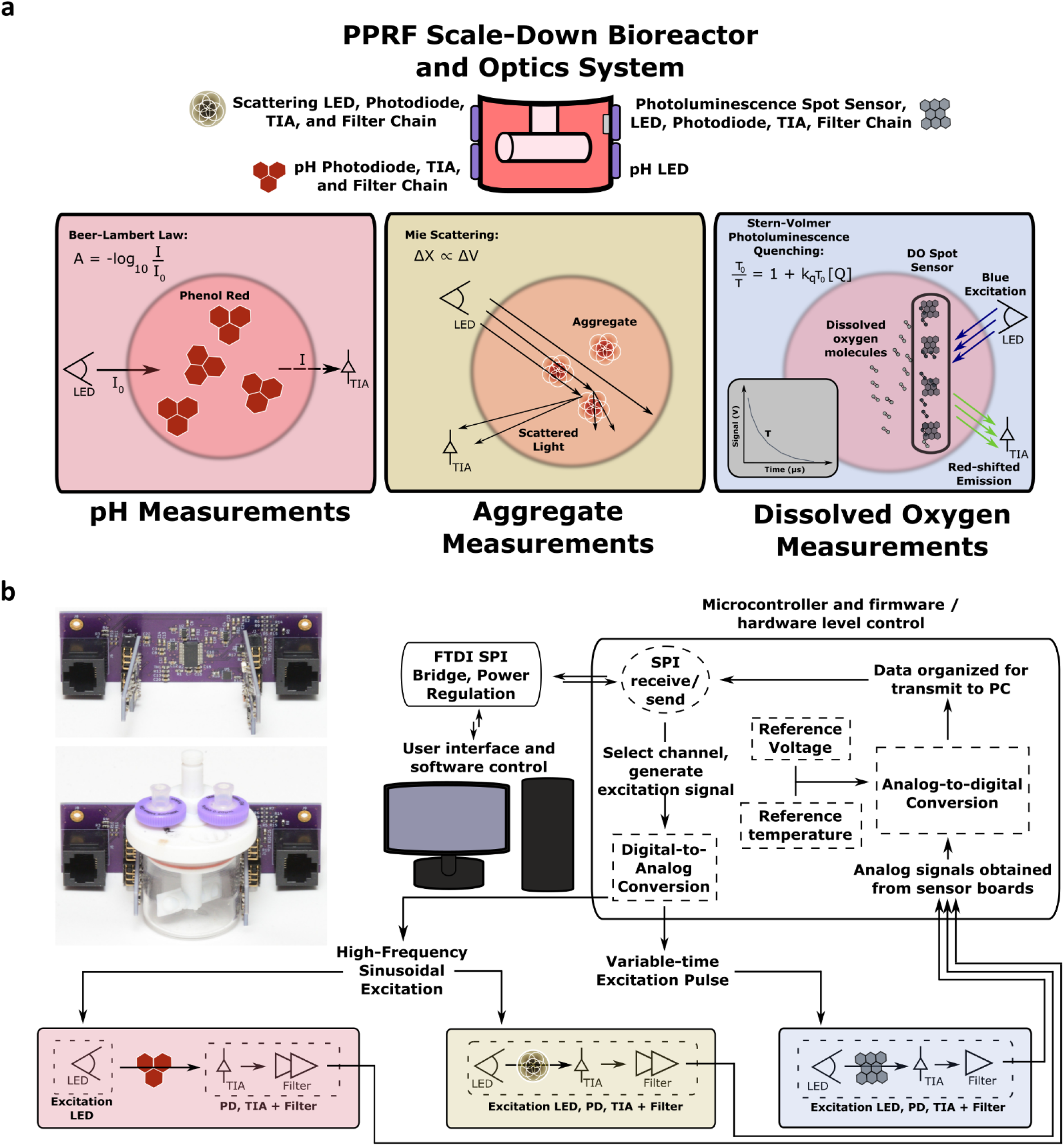
(a) Theoretical concepts governing function of the optoelectronics in measuring pH (left), cell density within the culture (center), and photoluminescence (right). (b) A picture of the system and its placement relative to the process vessel. A high-level schematic describes interactions between hardware in the system, from sensory components to digitization and data transfer.

## Methods

### Theoretical Principles Governing Measurements

Figure 2a provides a conceptual visualization of placement and interactions between optics and the culture vessel used in this study. The culture medium utilized for hiPSC expansion contains known concentrations of phenol-red, which in solution has absorbance characteristics that change deterministically with pH [24, 25]. The Beer-Lambert law was used to calculate pH from absorption of 550 nm excitation by phenol-red in the culture medium,

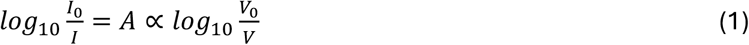

where transmitted intensity (I) is a function of the initial intensity (I0) and absorbance (A) of the interacting medium, measured voltages (V, V0) are proportional to optical intensity following photodetection. Calibration curves were derived from control measurements to characterize absorption relative to voltage as a function of pH in the medium. Scattering measurements were carried out using a 650 nm light-emitting diode (LED). The relationship between cell density of the population and optical scattering by cell aggregates in culture was modeled by linear regression and assessed by measuring voltage and analyzing relative to offline cell counts.

Oxygen-quenchable photoluminescent optochemical sensors were used to measure oxygen, where measurable lifetime of the active substrate is dependent on the concentration of dissolved oxygen (DO) present at the sensor interface [26, 27]. Optical excitation for photoluminescence phenomena was carried out using a 450 nm wavelength with a 550 nm longpass emission gel filter for detection. DO was calculated using the Stern-Volmer equation to describe changes in fluorescence lifetime (τ) proportional to oxygen concentration (Q),

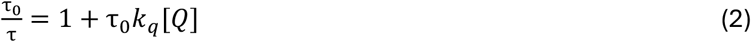

where k_q_ is the quenching reaction rate constant and τ0 is lifetime in the absence of oxygen.

### Hardware, Firmware, and Software Implementations

A PIC24FV16KM202 microcontroller (Microchip, USA) was utilized as the computing module for low-level control over system operation. Physical board layout and schematics are shown (Supp Figs 1, 2). The controller board was designed as a master interface to connect modular sensor boards each with their own regulation of power supply and independent acquisition pipeline. Current drivers were implemented on-board to generate sinusoidal excitations for absorbance and scattering measurements. A MOSFET driver was utilized for pulsed LED photoluminescence excitation. A thermistor was included to measure temperature dynamics of the system in-process, and a voltage reference included to assess electrical drift and supply voltage anomalies independent of the environment. An ADC acquisition speed of 80 kSps was used for pH, cell density, temperature, and reference voltage measurements, and 350 kSps to measure photoluminescence. All sampling periods stored 800 sequential 12-bit ADC acquisitions in memory per measurement. Onboard architecture was designed to facilitate daisy-chaining of controllers in series. This design permitted parallelization of data acquisition across multiple bioprocess vessels with minimal supporting hardware requirements.

Sensor boards included an LED specific to the wavelengths of interaction for each measurement. The pH channel included a separate board for the LED to be placed across the vessel from the sensor board. Excitation signals were generated at the microcontroller; sinusoidal current-driven signals were generated at 1 kHz for measurement of pH and cell density, and a 100 µs pulse was generated for photoluminescence measurements. The photoluminescence board was designed to include a tunable high-gain dc-biased transimpedance amplifier (TIA) followed by a fixed low-gain inverting amplifier stage. Sensor boards for pH and cell density were designed to include a tunable silicon photodiode (SiPD) TIA stage followed by a 1 kHz multiple feedback bandpass filter) with a −3 dB bandwidth of 0.97-1.03 kHz. An FT232H USB bridge (FTDI, Scotland) was used to interface a personal computer (PC) and the controller(s). Physical setup of the system and overall culture environment are shown in supplementary figure 3.

Firmware was written in Microchip C using MPLABX as a programming interface. Software was written in Python. The software runs in a loop, whereby the PC sends a command to the controller board(s) requesting measurement for a specific module in the system. After the data is captured digitally on the microcontroller, it is transferred to the PC and stored in a raw data file. This process continues for each module that is enabled in the software. After data for all enabled modules has been received, the device waits for a user-defined period, and the next iteration of the loop repeats. The program continues in this loop for a user-defined period or until interrupted.

### Data Processing Pipeline and Calculation of Metrics

Total acquisition times for absorbance and scattering were 12 ms each, and 0.25 ms for photoluminescence. A standardized pipeline for post-processing is shown (Fig 3). Prior to calculating amplitudes or applying an exponential fit, file size was reduced by applying 20x averaging to waveforms by ADC acquisition. Electrical drift was corrected by characterizing the relationship between system response and reference voltage drift. Temperature data allowed correction of temperature-associated deviations in reference and acquisition signals. These corrections were applied by characterizing system response as a function of the reference temperature. A protocol-gating algorithm was implemented utilizing temperature data to correct manual perturbations applied to each system during offline sampling (Fig 3, middle). The point at which a protocol deviation returned to within 0.2 °C of homeostasis was used as the constant of determination for protocol-gating. Offset correction was applied to the amplitude data to account for discrepancies in vessel placement relative to optical sensors following displacement during sampling (Fig 3, left). This was done by shifting all post-perturbation data to the mean of the last 100 data points prior, to maintain data continuity in post-processing. All data points during perturbation were set to this value. To estimate photoluminescent lifetime, a two-phase exponential was fitted to each exponential within the dataset to separate amplifier transients from the signal of interest (Fig 3, right).

**Figure 3.**
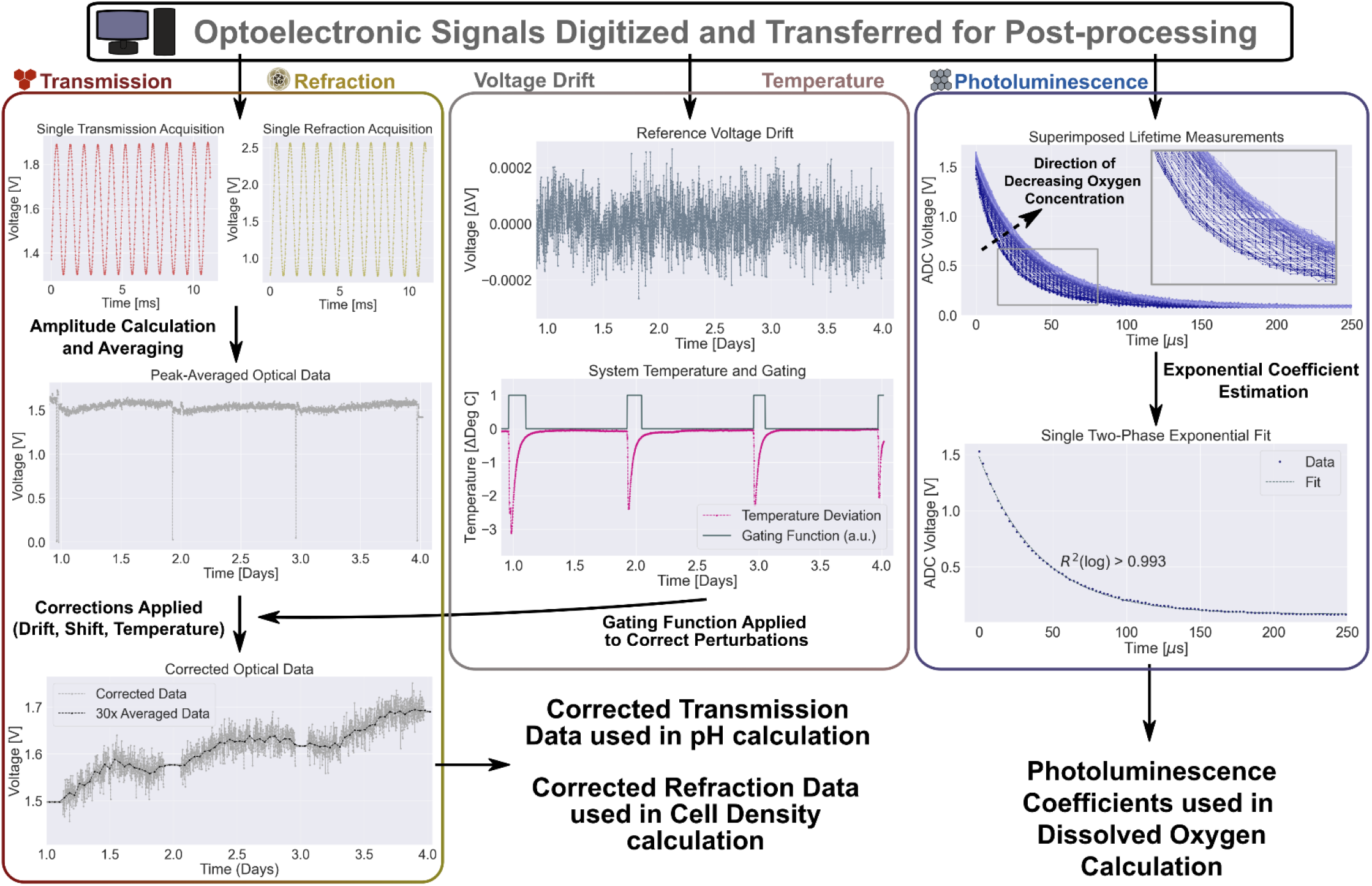
Post-processing pipeline for all channels following digitization and data transfer. Transmission and refraction data are averaged and plotted as sinusoidal amplitudes over time. Reference data (middle) is used to correct for both drift and misalignment following manual perturbation of the system during sampling. Note that the gating function is a binary variable of arbitrary unit, with values of 0 or 1. Exponential lifetime coefficients for the photoluminescent sensor are estimated by applying a two-phase exponential fit and isolating the coefficient relevant to the photoluminescence time (right).

pH calibration was performed under controlled experimental conditions, whereby an acidic solution containing equivalent concentration of phenol red to the culture medium was perturbed by controlled addition of sodium bicarbonate, and the pH response recorded by both the optoelectronics and an SBI ID Reader (Scientific Bioprocessing, Pennsylvania, USA). Absorbance was calculated where the intensity ratio was correlated to measured voltage ratio (Eq 1). Absorbance was plotted against measured pH in the linear region of interest to generate a calibration plot (Fig 4a).

**Figure 4.**
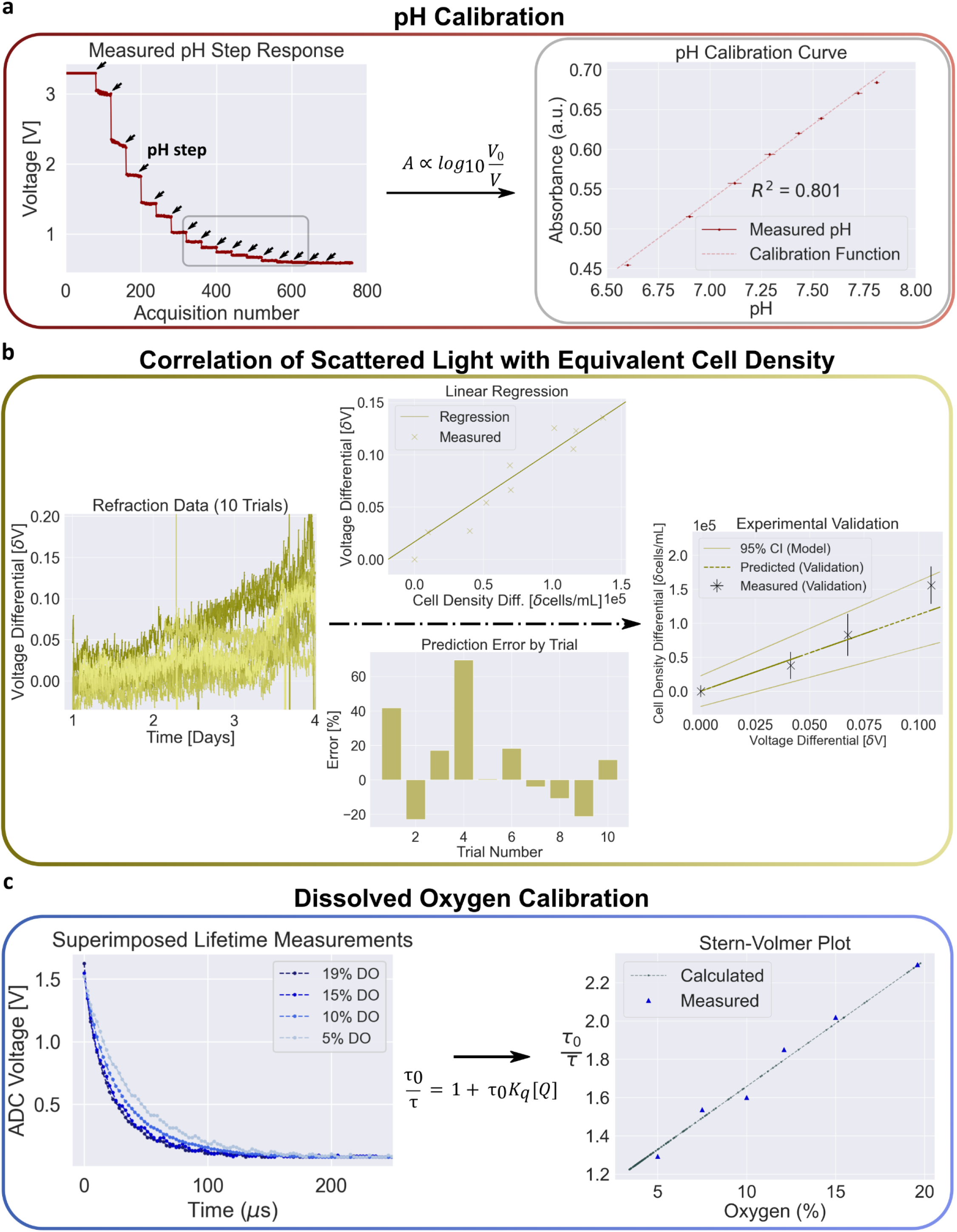
(a) Application of Beer-Lambert law to calculate pH following test measurements and resolve a calibration plot for linear region pH spanning 6.6-7.8. (b) Implementation of a linear regression model to estimate correlation of cell density and signal deltas for trial experiments. Error is shown as the percentage difference from the control measurement using the training data. The model was applied to a validation culture and plotted relative to the confidence intervals (CI) to determine utility of the model. (c) Measurements for manual acquired photoluminescence during nitrogen purge experiments. Manual recordings against calculated Stern-Volmer relation for the region of oxygen under study (3.5-19.7%).

A linear regression strategy was chosen for its performance across repeated experiments between measured growth and observed voltage response at the refraction channel. To calculate cell density, a linear empirical model was applied to 10 datasets from hiPSC expansion experiments to correlate scattering with cell density relative to offline measurements at the end of each run (Fig 4b). Model performance was compared to discrete control measurements taken at the end of each experiment, and absolute error assessed as the percentage difference between predicted and actual values. The model was then applied to a test culture, where offline measurements were taken daily. Continuous predicted and discrete measured results were compared alongside confidence intervals calculated from the training set.

To determine quenching rate constant and baseline lifetime in the absence of a quenching species, calibration experiments were performed for each batch of dissolved oxygen sensors used. Sterilized vessels containing culture medium were fitted with an adherent PSt3-NAU chemosensor spot (PreSens, Germany) at the interior glass interface of the reactors under sterile conditions. Vessels were placed in a hypoxia chamber (Supp Fig 2b). Environmental oxygen was purged and monitored using a ProOx model P110 compact O2 controller, gas inlet, and peripheral probe (BioSpherix, New York, USA). Compressed nitrogen was used to purge oxygen levels inside the hypoxia chamber from 19.7% ambient down to a setpoint of 3.5%. Intermediate environmental measurements were recorded from the oxygen controller. Stern-Volmer relation was used to characterize optochemical sensor response to oxygen (Fig 4c).

### Scale-down hiPSC Expansion Bioprocess Protocols

This work was carried out under ethics protocol #REB14-1914, approved by the University of Calgary Conjoint Health Research Ethics Board (CHREB). Expansion protocols for hiPSCs have been developed and rigorously optimized at scales larger than 50 mL. Protocols for scale-down experiments were carried out as described previously for larger vessels, with static controls [28, 29]. Notable differences to facilitate optimized growth under scale-down conditions are included herein. hiPSC cell line, designated PGPC14 (XX-chromosome) at passage 19 was kindly provided by James Ellis’ lab at University of Toronto and used for this study [28, 29].

Following a single static passage, aggregates were preformed in 12-well tissue-culture treated plastic plates in 1 mL mTeSR1 (cat#85850, StemCell Technologies, Vancouver, Canada) supplemented with 10 µM Y-27632 (cat#72304, StemCell Technologies) for 24 hours prior to inoculation in horizontal blade scale-down reactors. Agitation was controlled at 150 RPM using a bioWiggler programmable stirrer platform (VWR, Pennsylvania, USA). A dynamic inoculation density of 6.8×10^4^ cells/mL as preformed aggregates was used. Cultures were carried out over 4 days following static passage (including preformation of aggregates).

Environmental chamber oxygen was monitored using a ProOx model P110 compact O2 controller and peripheral probe. Carbon dioxide was controlled via a ProCO2 model 120 controller and peripheral probe (BioSpherix, New York, USA). Cultures were maintained at 37 ° C, >88% humidity, and 5% ambient CO2 under incubation. A reservoir of H2O was retained in the hypoxia chamber to minimize liquid medium evaporation. Vessels were removed from the optoelectronic frame for offline sampling. For offline images and counts, 1.5 mL samples were taken with medium replacement. Offline counts were measured using a NucleoCounter and processed in NucleoView (ChemoMetec, Denmark). Figure 5 provides a typical experimental timeline for cultivation of hiPSCs in the scale-down system, coupled with online monitoring results obtained during culture.

**Figure 5.**
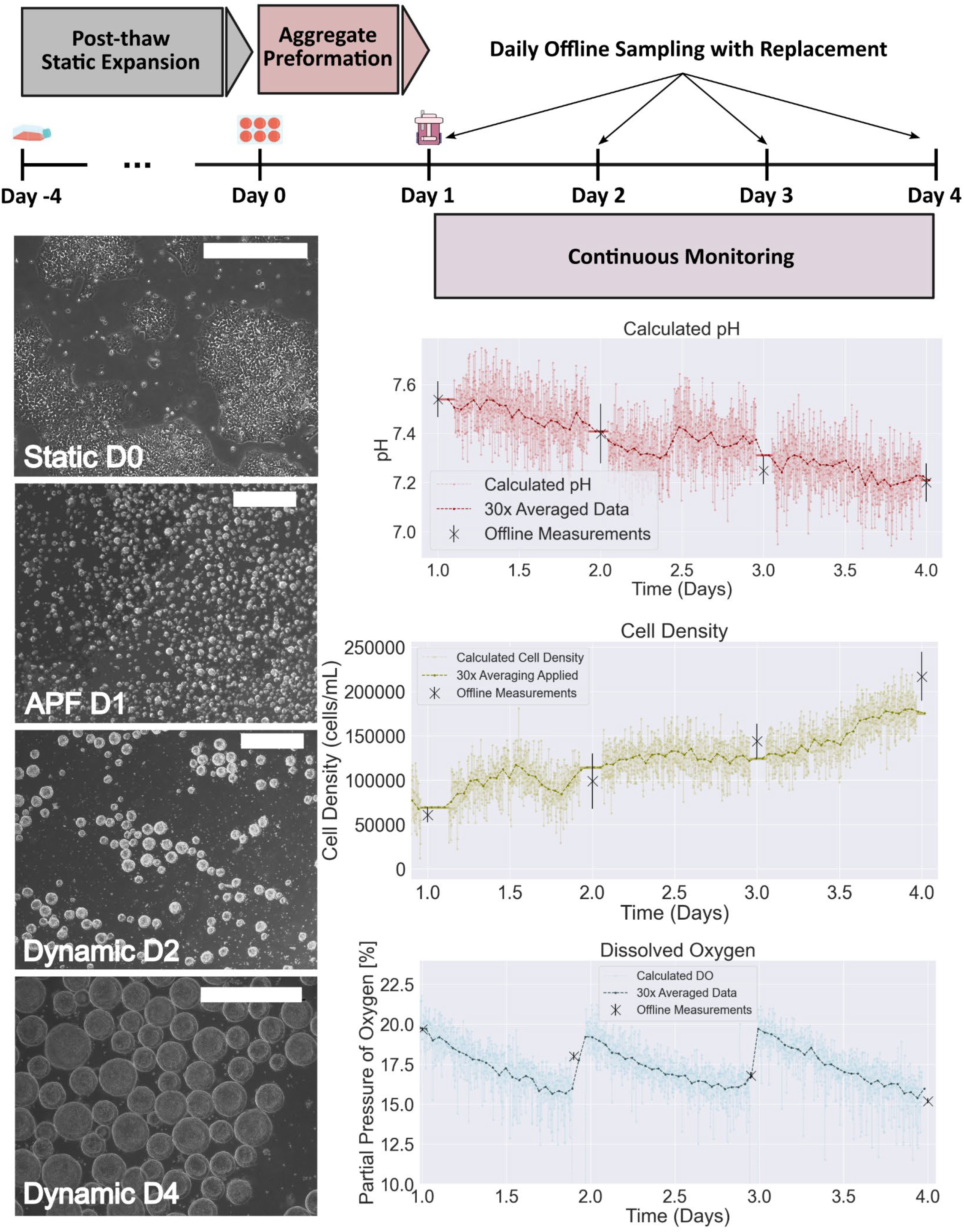
(top) Bioprocess timeline for hiPSC expansion, acquisition of online data, and sampling for offline measurements. (left) morphological characteristics following four-day static expansion, 24-hour aggregate preformation (APF), and three-day dynamic expansion. (right) Calculated parameters from instrumentation data for pH, cell density, and dissolved oxygen throughout dynamic expansion. Scale-bars are 500 µm.

### Application Study – *in vitro* Oxygen Deprivation

To test the system, data from hypoxic and normoxic culture conditions were acquired during hiPSC cultivation over a single passage (Fig 6). Under hypoxia, environmental oxygen was purged from the system to a setpoint of 3.5% DO several hours following inoculation. Note that no offline sampling was performed prior to harvest on day 4. Cell-normalized relative oxygen consumption rate (OCR) in the system were calculated as:

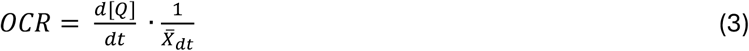

where [Q] is the oxygen concentration at time t, and 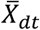 is mean cell concentration across the time interval over which OCR is calculated. Extracellular acidification rate (ECAR) was calculated as the cell-normalized change in pH of the process environment over time:

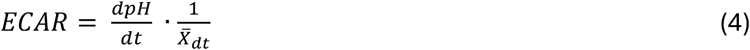

**Figure 6.**
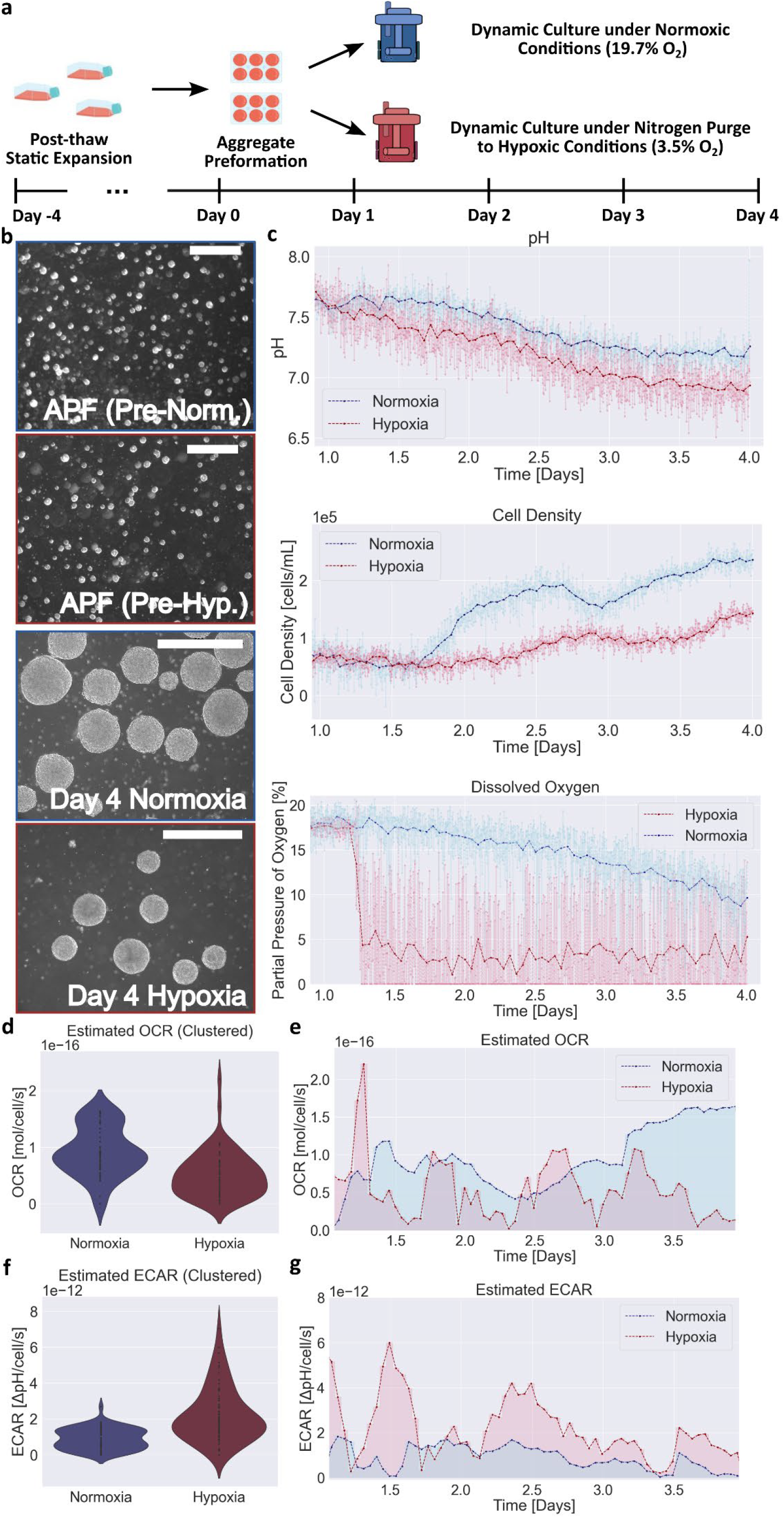
Characterization of metabolic phenotype in iPSCs cultured under normoxia and hypoxia in a scale-down reactor environment. (a) Timeline for the protocols used in oxygenation experiments is shown. (b) Morphological characteristics following aggregate preformation and at day 4 following dynamic culture. (c) Continuous data acquired over experiment for pH, dissolved oxygen, and cell density. (d) Calculated OCR, aggregated across the experiment. (e) Temporally resolved OCR calculations. (f) Calculated ECAR, aggregated across the experiment. (g) Temporally resolved ECAR calculations. All scalebars are 500 µm.

OCR and ECAR were calculated using the above equations. To correct for contributions from the experimental setup to oxygen losses in the system during nitrogen purge, vessel oxygen was measured in a control condition in the absence of cells prior to equilibrium. The oxygen transfer rate out of the system during purge was calculated using the control data and subtracted from the calculated OCR under hypoxia prior to observed equilibrium. Processed data were averaged over ±2.5 hours at each time-point used in calculation of OCR and ECAR. To further validate instrumentation capability in capturing surrogate measures of metabolic function *in vitro*, additional cultures were carried out over 4 days under normoxic or hypoxic condition (Supp Fig 4).

### Statistics

Three technical replicates were taken for in vitro offline measurements. Online measurement data were plotted as 20x averaged and 600x averaged data for a single monitored vessel. This corresponded to 2-minute or 1-hour windows, respectively. Offline comparison data were plotted as mean ± standard deviation. Coefficient of determination (R2) was used to represent proportion of variance in the dependent variable explained by linear fits for pH and cell density calculation. R^2^(log) was used to represent variance for exponential fits. Absolute error was reported as a metric of linear regression performance for cell density calculations. Confidence intervals were calculated from a two-sided t-test statistic obtained from the linear regression training dataset.

## Results

### Optoelectronic Instrumentation

The system was sufficiently optically isolated to prevent ambient optical noise in an incubator environment. The primary source of perturbations to signals were temperature and manual sampling. Voltage drift (DC) over time across the entirety of experiments observed in the reference channel was less than 1 mV for all measurements. Temperature variance of <0.2 °C was exhibited *in vitro* at equilibrium, with deviations up to 5.0 °C during culture protocol perturbations. Approximate power consumption during sampling was ~0.2 W per system, and <0.04 W during transmit and idle. Sampling time accounted for 8% of total active time (24.25 ms per 300 ms). The system was activated to acquire and transmit data once every 6 s.

### Processing Pipeline and Modeling

Manual perturbations were successfully corrected by utilizing reference temperature and voltage. Correction for offset in the system was significant in improving the quality of obtained data in reflecting dynamics within the culture vessel. While temperature correction was relevant to the measurement data in the event of perturbation, no significant improvement was made by temperature-correcting the raw datasets following protocol-gating and shift-correction.

Resolutions for each channel were determined from the standard deviation of post-processed data following 600x averaging, based on quantification of calibration data and modeling results (Fig 4). A standard deviation of 0.05 for pH within the linear region of phenol-red absorption was observed – It is important to note that absorbance corrections due to variance in temperature were explored, but these corrections did not improve resolution of the system beyond the variance introduced by optoelectronic noise during measurement. For univariate linear modeling of voltage in response to cell density, R^2^ > 0.57 was obtained. Root mean squared error (RMSE) of 22.94% was obtained across cell density estimations. As a single agitation rate was used for these experiments, no consideration was made for correcting signals relative to aggregate distributions within the culture, or in characterizing biological and protocol-associated variance relative to error observed in the measurements. With respect to dissolved oxygen, a standard deviation of 0.4% was observed – No corrections were made for sensor-to-sensor variability outside of batch calibration. R^2^ > 0.993 was obtained for two-phase exponential fits relative to photoluminescence measurements.

### Scale-Down hiPSC Bioprocess Outcomes

Completion of the scale-down protocol under normoxic conditions resulted in expected growth and proliferation (Fig 5). Morphological outcomes for static colonies and aggregates were normal, and a definitive increase in aggregate diameter was observed over time (Fig 5, left). Offline measurements of cell density were negatively correlated with both pH and ambient oxygen and corroborated by online data acquired over experimental time-course (Fig 5, right). Online pH, dissolved oxygen, and cell-density measurements followed expected trajectories.

### Application Study Results – *in vitro* Oxygen Deprivation

Following oxygen deprivation, cell growth was markedly reduced under hypoxic conditions (Fig 6b, c). Images taken on day 4 showed a smaller number of aggregates with reduced diameter and slight morphological differences under hypoxia, and a smaller proportion of single cells in suspension (a result of aggregate disruption during sampling). pH of both systems was comparable, and oxygen measurements were observed to reflect gas conditions in the vessel. Cell-normalized estimates for OCR and ECAR show a clear differential between normoxic and hypoxic conditions towards day 4 of culture, with dynamic behavior observed throughout the experiment time course (Fig 6d-f). Further, follow-up experiments confirm consistent relative measurements and metabolic activity between conditions, reinforcing confidence in utilizing pH, aggregate density, and oxygen measurements to calculate oxygen consumption and extracellular acidification dynamics. (Supp Fig 4a, 4b).

## Discussion

We have presented herein our characterization and implementation of an approach to monitor surrogate measures of metabolic phenotype. We evaluated the functionality and feasibility of using our optoelectronic system for continuous data collection of key bioprocess parameters. Our instrumentation work demonstrated capable performance of a low-cost open-source system with minimal infrastructural requirements. The materials cost of the instrumentation utilized in this work is approximately $60 CAD, making it similar in cost to a single spot sensor for optical measurement of dissolved oxygen. These characteristics make our instrumentation approach more financially accessible for parallelizing experimental setups in an academic setting relative to commercially available systems such as the Ambr15 from Sartorius (costing upwards of 50,000 USD), although this must be tempered by constraints on system resolution. It should also be noted that many medium formulations are absent of phenol-red. Transition of pH measurement away from phenol-red absorbance to optical spot sensors would greatly improve universality of the system, and is entirely feasible under the current instrumentation constraints.

This work has also provided confidence in our current endeavors to couple online measurements with omics data for translation to high-dimensionality phenotypic modeling in hiPSC bioprocesses. Of most interest to our group is to assess the extent to which we can couple surrogate measures of phenotype to more complex activity in the cell population in conjunction with time-series multi-omics datasets and hydrodynamic simulations. Continuation of this work will provide further insights into dynamic cell state as a result of artificial environmental setpoints that are meant to recapitulate the complex embryonic niche in stasis [8, 30–37].It should be noted that significant disruption to measurement is observed during manual culture-protocol-associated perturbations. This is due to the tolerance of the frame and optoelectronics positioning relative to the vessel – displacement of up to 2 mm results in significant alteration of measured voltage given the curved optical interface of the culture vessel itself. The reliability of the system could be improved through adjustments to both the mechanical implementation and optical design, so as to eliminate the need for corrections to these perturbations, or a software approach to correct or eliminate signal contributions from manual perturbations in real-time.

Aggregate density, size, and spatial distribution within the bioprocess are expected to have significant effects on our ability to measure cell density – these are all variables that are inextricably tied to the hydrodynamics of the culture system [32]. Our model could be improved by capturing signal characteristics as a function of aggregate distributions, density, and spatial dynamics within the process vessel. As the system geometry and agitation dynamics are changed, these variables are altered leading to changes in the interaction of optics and biological constituents. Further, significant disruption is observed during manual culture-protocol-associated perturbations. This is due to the tolerance of the frame and optoelectronics positioning relative to the vessel – displacement of up to 2 mm results in significant alteration of measured voltage given the curved optical interface of the culture vessel itself. The reliability of the system could be improved through adjustments to both the mechanical implementation and optical design, so as to eliminate the need for corrections to these perturbations.

As aggregate distributions are largely affected by hydrodynamic conditions within the vessel, alterations to agitation rate or adaptation to another impeller and vessel geometry will require re-modeling of the signal characteristics relative to cell growth, or inclusion of variables to explain optical perturbations arising from these interactions. The authors are confident that reported values for scattering model accuracy are a stepping-stone for the development of more advanced multivariate implementations, and that remarkable improvements can be made across this approach, including through the implementation of more advanced optics. While several technologies have been applied to measure cell densities or surrogates thereof to deduce cellular dynamics in culture, we have not observed in literature non-invasive approaches applied to measuring optical scattering by aggregates throughout expansion in hiPSC bioprocessing [38–40]. Without continuous reference data against our measurements, and with a lack of exhaustive time-series hiPSC culture data in literature, it is difficult to surmise the root cause of plateauing or oscillatory dynamics observed in our data throughout the experiments. Whether these signal responses are the result of environmental artifacts or cell population dynamics is unclear and warrants further studies.

With regards to OCR, purging oxygen from the environment through the introduction of pressurized nitrogen results in oxygen losses across multiple interfaces. These are incurred from the vessel itself due to increased environmental pressure coupled with reduced partial pressure of oxygen at the gas-liquid interface, and through latent losses of oxygen at the junctions of the hypoxia chamber. Effectively characterizing the non-cell associated losses of oxygen from within the vessel was successful in accounting for a large proportion of the oxygen losses observed during purge, though corrected oxygen consumption rate was several times higher in the hypoxic condition during purge. Whether this outcome is a result of incomplete characterization of the non-cell mediated losses or is representative of an acute hyperoxia response by the cell population is unknown and requires further investigation. It must be stated that without characterization of the mass transfer coefficients associated with the environmental dynamics of the culture system, OCR calculations are limited in accuracy. Indeed, while the values calculated herein are physiologically viable, they tend towards the upper limit of OCR reported in literature for hiPSCs, suggesting that losses in the chamber system may potentially inflate the estimates. Further, specific oxygen uptake rates (OUR) of the cells cannot be calculated [41, 42]. However, these measurements provide insights into oxygen characteristics of the process environment and subsequently the artificial niche [43–46]. Further, given the relative nature of the measurements in comparing population conditions, the outcome is meaningful in differentially assessing environmental impact on the cell population within the niche.

The authors are confident that the ability to capture differential behavior between conditions with high temporal resolution provides a foundation to inform complex in-process phenotypic behavior in combination with multi-omics characterization. Further, given that the geometry of the vessels utilized in this research are conducive to empirical modeling and simulation approaches to scale-up, the authors are confident that findings observed at this scale are translatable to commercial manufacturing scales [47]. While our results are promising, there is ample room for improvement in the quality of the approach to collecting parallelized parametric data across biological replicates under scale-down conditions or applying these perspectives to clinical scale manufacturing infrastructure for optimized vessel geometries. We believe that adapting monitoring and characterization strategies to more thoughtfully capture aspects of cell phenotype will improve visibility of cell state dynamics that may be significant in tailoring process pipelines to therapeutic derivatives [48, 49].

## Conclusion

Our group developed, implemented, and validated a modular open-source continuous monitoring system for use in characterizing surrogate metabolic metrics of phenotype in scale-down vessels. We demonstrated the ability of the system to utilize key process parameters in establishing a surrogate estimate of metabolic phenotype. Metabolic remodeling of the cell population constitutes important indicators of cell state within pluripotency and this data provides us with additional insight to explore the trajectory of the cell population in conjunction with higher resolution cell characterization data. Further, we can apply these principles to the capture of more complex cell state dynamics by implementing sensing modalities to measure metabolic intermediates within the bioprocess. The results of this study support this approach for applications in modeling in vitro phenotype dynamics in conjunction with offline multi-omics characterization datasets. Tools such as the one developed for this study provide researchers with open-source, affordable and adaptable approaches outside of conventional scale bio-manufacturing. While a plethora of limitations and boundaries exist, researchers around the globe are intent on overcoming these obstacles in unique and impactful ways to shape the future of regenerative medicine in healthcare.

## Acknowledgements

We gratefully acknowledge the Canadian Institutes for Health Research (CIHR), the Natural Sciences and Engineering Research Council of Canada (NSERC), and CMC Microsystems for supporting this work.

## Author Contributions

All authors conceived the approach to monitoring a surrogate measure of metabolic phenotype *in vitro*. J.C. designed and tested all instrumentation under the mentorship of K.M. J.C. carried out all hiPSC culture experimental work under the mentorship of M.K. All software was developed by J.C. J.C. prepared the manuscript and all visualization. All authors contributed to edits and revised the manuscript.

## Data Availability Statement

All material available upon request to the corresponding author. Consolidated data and code are publicly available at: https://github.com/JxColter/Optoelectronics-Colter-J.-et-al.-2025/

## Additional Information

The authors declare that they have no competing or conflicting interests.

## References

[1] A. Yilmaz & N. Benvenisty, “Defining human pluripotency”, Cell Stem Cell, vol. 25, no. 1, pp. 9–22, July 2019, 10.1016/j.stem.2019.06.010

[2] M. Patterson et al., “Defining the nature of human pluripotent stem cell progeny”, Cell Research, vol. 22, pp. 178–193, Aug 2011, 10.1038/cr.2011.133

[3] Y. Shi et al., “Induced pluripotent stem cell technology: a decade of progress,” Nature Reviews Drug Discovery, vol. 16, pp. 115–130, Dec 2016, 10.1038/nrd.2016.245

[4] M.A. Glicksman, “Induced pluripotent stem cells: The most versatile source for cell therapy”, Clinical Therapeutics, vol. 40, no. 7, pp.1060–1065, July 2018, 10.1016/j.clinthera.2018.06.004

[5] M. Madrid et al., “Autologous induced pluripotent stem-cell based cell therapies: Promise, progress, and challenges,” Current Protocols, vol. 1, no. e88, pp. 1–25, Mar 2021, 10.1002/cpz1.88

[6] J. Deinsberger, D. Reisinger, & B. Weber, “Global trends in clinical trials involving pluripotent stem cells: a systematic multi-database analysis,” npj Regen Med, vol. 5, no. 15, pp. 1–13, Sep 2020, doi: 10.1038/s41536-020-00100-4

[7] J.Y. Kim et al., “Review of the current trends in clinical trials involving pluripotent stem cells,” Stem Cell Rev and Rep, vol. 18, pp. 142–154, Sep 2021, 10.1007/s12015-021-10262-3

[8] Colter J. et al., “Induced pluripotency in the context of stem cell expansion bioprocess development, optimization, and manufacturing: a roadmap to the clinic,” npj Regen Med, vol. 6, no. 72, pp. 1–9, Nov 2021, 10.1038/s41536-021-00183-7

[9] S. Kobold et al., “History and current status of clinical studies using pluripotent stem cells,” Stem Cell Reports, vol. 8, pp. 1–7, Jul 2023, 10.1016/j.stemcr.2023.03.005

[10] R. Lister et al., “Hotspots of aberrant epigenomic reprogramming in human induced pluripotent stem cells,” Nature, vol. 471, pp. 68–73, Feb 2011, 10.1038/nature09798

[11] S. Efrat, “Epigenetic memory: Lessons from iPS cells derived from human β cells,” Front. Endocrinol., vol. 11, pp. 1–5, Jan 2021, 10.3389/fendo.2020.614234

[12] S. Yamanaka, “Pluripotent stem cell-based cell therapy – Promise and challenges,” Cell Stem Cell, vol. 27, pp. 523–531, Oct 2020, 10.1016/j.stem.2020.09.014

[13] X. Liu et al., “Chromosomal aberration arises during somatic reprogramming to pluripotent stem cells,” Cell Div, vol. 15, no. 12, pp. 1–13, Oct 2020, 10.1186/s13008-020-00068-z

[14] T. Iida et al., “Whole-genome DNA methylation analyses revealed epigenetic instability in tumorigenic human iPSC cell-derived neural stem/progenitor cells,” Stem Cells, vol. 35, pp. 1316–1327, Jan 2017, 10.1002/stem.2581

[15] T.S. Khoo et al., “Retention of somatic memory associated with cell identity, age, and metabolism in induced pluripotent stem (iPS) cells reprogramming,” Stem Cell Rev and Rep, vol. 16, pp. 251–261, Feb 2020, 10.1007/s12015-020-09956-x

[16] V. Volpato & C. Webber, “Addressing variability in iPSC-derived models of human disease: Guidelines to promote reproducibility,” Dis. Model. & Mech., vol. 13, pp. 1–9, Jan 2020, https://doi.org/1242/dmm.042317

[17] H. Kilpinen et al., “Common genetic variation drives molecular heterogeneity in human iPSCs”, Nature, vol. 546, pp. 370–375, May 2017, 10.1038/nature22403

[18] A. Kurtz et al., “Linking scattered stem cell-based data to advance therapeutic development,” Trends in Molecular Medicine, vol. 25, No. 1, pp. 8–19, Jan 2019, 10.1016/j.molmed.2018.10.008

[19] Jenkins, M. et al., “Patient-specific hiPSC bioprocessing for drug screening: Bioprocess economics and optimization,” Biochemical Engineering Journal, vol. 108, pp. 84–97, Apr 2016, 10.1016/j.bej.2015.09.024

[20] K. Nishimura, A. Fukuda, & K. Hisatake, “Mechanisms of the metabolic shift during somatic cell reprogramming,” Int. J. Mol. Sci., vol. 20, No. 2254, pp. 1–16, May 2019, 10.3390/ijms20092254

[21] J. Prieto et al., “Mitochondrial dynamics and metabolism in induced pluripotency,” Experimental Gerontology, vol. 133, pp.2–19, Feb 2020, 10.1016/j.exger.2020.110870

[22] S. Burr, A. Caldwell, M. Chong et al., “Oxygen gradients can determine epigenetic asymmetry and cellular differentiation via differential regulation of Tet activity in embryonic stem cells,” Nucleic Acids Research vol. 46, No. 3, Nov 2017. 10.1093/nar/gkx1197

[23] M. Belli, P. Rinaudo, M.G. Palmerini et al., “PreImplantation Mouse Embryos Cultured In Vitro under Different Oxygen Concentrations Show Altered Ultrastructures,” Int. J. Environ. Res. Public Health vol. 17, No. 10, May 2020. 10.3390/ijerph17103384

[24] J. Michl et al., “Evidence-based guidelines for controlling pH in mammalian live-cell culture systems, Commun. Biol., vol. 2, No. 144, Apr 2019, 10.1038/s42003-019-0393-7

[25] L. Rovati et al., “Plastic Optical Fiber pH Sensor using a Sol-Gel Sensing Matrix,” Fiber Optic Sensors InTech, Feb 2012, 10.5772/26517

[26] J. Yang et al., “Real-time monitoring of dissolved oxygen with inherent oxygen-sensitive centers in metal-organic frameworks,” Chem. Mater., vol. 28, no. 8, pp. 2652–2658, Apr 2016, 10.1021/acs.chemmater.6b00016

[27] T. Burger et al., “Porphyrin based metal-organic frameworks: highly sensitive materials for optical sensing of oxygen in gas phase,” J. Mater. Chem. C, vol. 9, pp. 17099–17112, Nov 2021, 10.1039/D1TC03735H

[28] B.S. Borys et al., “Optimized serial expansion of human induced pluripotent stem cells using low-density inoculation to generate clinically relevant quantities in vertical-wheel bioreactors,” Stem Cells Translational Medicine, vol. 9, no. 9, pp. 1036–1052, Sep 2020, 10.1002/sctm.19-0406

[29] B.S. Borys et al., “Overcoming bioprocess bottlenecks in the large-scale expansion of high-quality hiPSC aggregates in vertical-wheel stirred suspension bioreactors,” Stem Cell Research & Therapy, vol. 12, no. 55, pp. 1–19, Jan 2021, 10.1186/s13287-020-02109-4

[30] M.S. Reuter et al., “The personal genome project Canada: Findings from whole genome sequences of the inaugural 56 participants,” CMAJ, vol. 190, no. 5, pp. e126–e130, Feb 2018. 10.1503/cmaj.171151

[31] M.R. Hildebrandt et al., “Precision health resource of control iPSC lines for multilineage differentiation,” vol. 13, no. 6, pp. 1126–1141, Dec 2019, 10.1016/j.stemcr.2019.11.003

[32] E. Csaszar et al., “Process evolution in cell and gene therapy from discovery to commercialization,” Can. J. Chem. Eng., vol. 99 (11), pp. 2517–2524, Nov 2021, 10.1002/cjce.24141

[33] A. Riviera-Ordaz et al., “Critical analysis of cGMP large-scale expansion process in bioreactors of human induced pluripotent stem cells in the framework of quality by design,” BioDrugs, vol. 35, pp. 693–714, Nov 2021, 10.1007/s40259-021-00503-9

[34] T. Dang et al., “Computational fluid dynamic characterization of vertical-wheel bioreactors used for effective scale-up of human induced pluripotent stem cell aggregate culture,” Can. J. Chem. Eng., vol. 99, pp. 2536–2553, Jul 2021, 10.1002/cjce.24253

[35] A. Polanco, B. Kuang & S. Yoon, “Bioprocess technologies that preserve the quality of iPSCs,” Trends in Biotechnology, vol. 38, no. 10, pp.1128 – 1140, Oct 2020, 10.1016/j.tibtech.2020.03.006

[36] F. Manstein et al., “High density bioprocessing of human pluripotent stem cells by metabolic control and in silico modeling,” Stem Cells Translational Medicine, vol. 10, no. 7, pp. 1063–1080, Jul 2021, 10.1002/sctm.20-0453

[37] P. Moutsatsou et al., “Automation in cell and gene therapy manufacturing: from past to future,” Biotechnology Letters, vol. 41, pp. 1245–1253, Sep 2019, 10.1007/s10529-019-02732-z

[38] K.M. Fridley, M.A. Kinney & T.C. McDevitt, “Hydrodynamic modulation of pluripotent stem cells,” Stem Cell Research & Therapy, vol. 3, no. 45, pp. 1–9, Nov 2012, 10.1186/scrt136

[39] C. Kropp, et al., “Impact of feeding strategies on the scalable expansion of human pluripotent stem cells in single-use stirred tank bioreactors,” Stem Cells Translational Medicine, vol. 5, no. 10, pp. 1289–1301, Oct 2016, 10.5966/sctm.2015-0253

[40] X. Liu, et al., “The uniformity and stability of cellular mass density in mammalian culture,” Front. Cell Dev. Biol. vol. 10, Oct 2022, 10.3389/fcell.2022.1017499

[41] G.E. Neurohr & A. Amon, “Relevance and regulation of cell density,” Trends in Cell Biology vol. 30, no. 3, Mar 2020, 10.1016/j.tcb.2019.12.006

[42] J. Carvell & J.E. Dowd, “Online measurements and control of viable cell density in cell culture manufacturing processes using radio-frequency impedance,” Cytotechnology vol. 50, Jun 2006, 10.1007/s10616-005-3974-x

[43] M. H. Gehlan, “The centenary of the Stern-Volmer equation of fluorescence quenching: From the single line plot to the SV quenching map,” J. Photochem. Photobio. C: Photochem. Rev. vol. 42, Mar 2020, 10.1016/j.jphotochemrev.2019.100338

[44] M. Pappenreiter et al., “Oxygen uptake rate soft-sensing via dynamic kLa computation: Cell volume and metabolic transition prediction in mammalian bioprocesses,” Front. Bioeng. Biotechnol. vol. 7, no. 195, pp. 1–16, Aug 2019, 10.3389/fbioe.2019.00195

[45] E. Tanumihardja et al., “Measuring both pH and O2 with a single on-chip sensor in cultures of human pluripotent stem cell-derived cardiomyocytes to track induced changes in cellular metabolism,” ACS Sensors, vol. 6, no. 1, Jan 2021, 10.1021/acssensors.0c02282

[46] A. Super et al., “Real-time monitoring of specific oxygen uptake rates of embryonic stem cells in a microfluidic culture device,” J. Biotechnol. vol. 11, no. 9, Sep 2016, 10.1002/biot.201500479

[47] Borys, B.S., Roberts, E.L., Le, A. et al. “Scale-up of embryonic stem cell aggregate stirred suspension bioreactor culture enabled by computational fluid dynamics modeling,” Biochemical Engineering Journal vol. 133, May 2018, 10.1016/J.BEJ.2018.02.005

[48] Y. Kim et al., “Scalable manufacturing of solderable and stretchable physiologic sensing systems,” Adv. Mat., vol. 29, pp. 1701312 – 1701323, Aug 2017, 10.1002/adma.201701312

[49] M. Goker et al., “Bioprocess monitoring by bio-sensor based technologies,” Clinical, Food, and Beyond, pp. 259–285, 10.1016/B978-0-12-818592-6.00010-4

