## Supplemental Figures for "Online characterization of surrogate metrics for metabolic phenotype in human induced pluripotent stem cell bioprocessing"

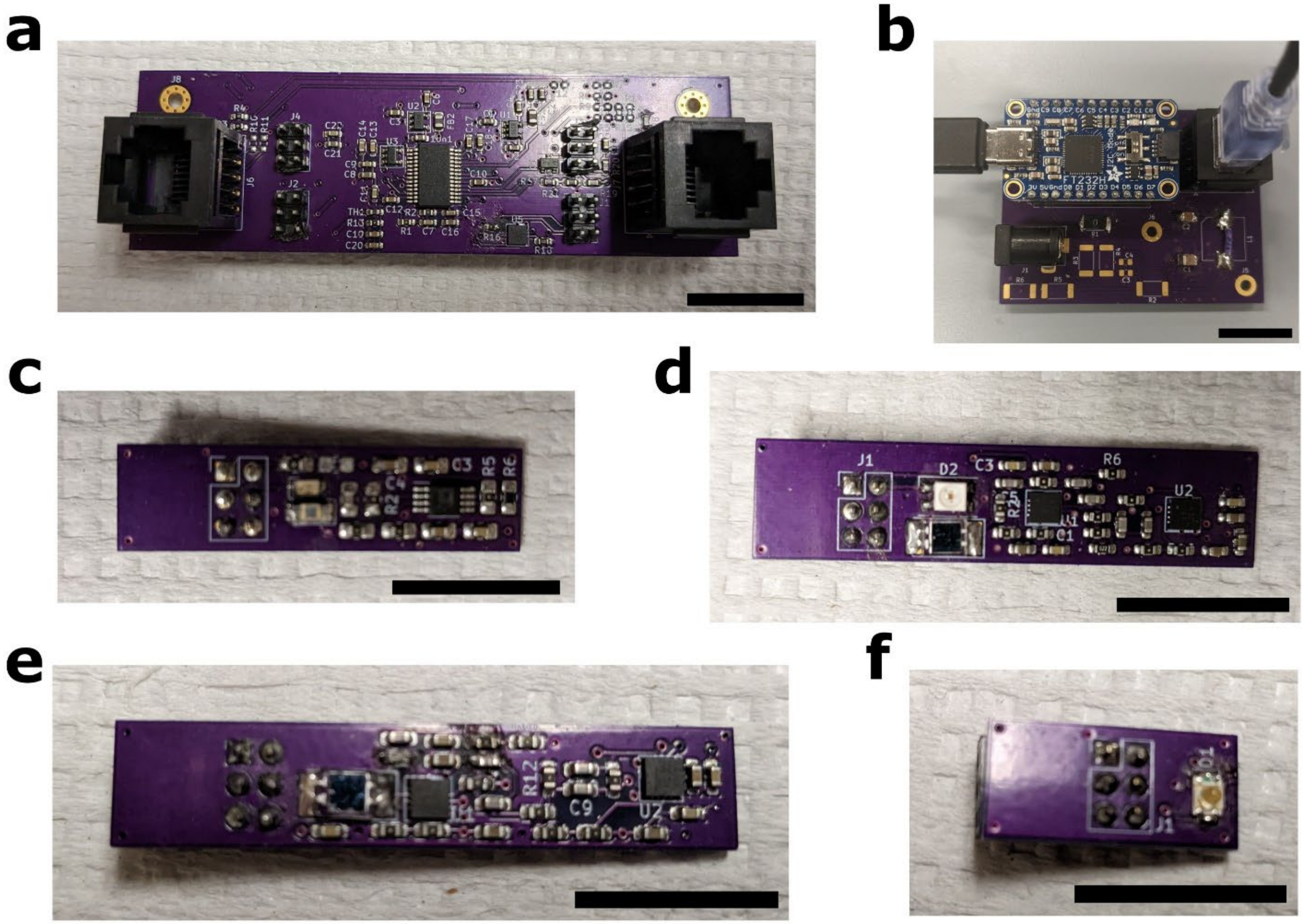


Supplementary Figure 1. Individual components making up the optoelectronic instrumentation. (a) Control board, incorporating the microcontroller, onboard voltage regulation, temperature sensor, digital/analog signal routing, and drivers for optical excitation. (b) Interface board, incorporating an FTDI SPI interface, signal routing and additional power regulation capabilities. (c) Photoluminescence instrumentation. (d) Refraction instrumentation. (e) Transmission emission instrumentation. (f) Transmission LED board. Scale-bars are 2.5 cm.

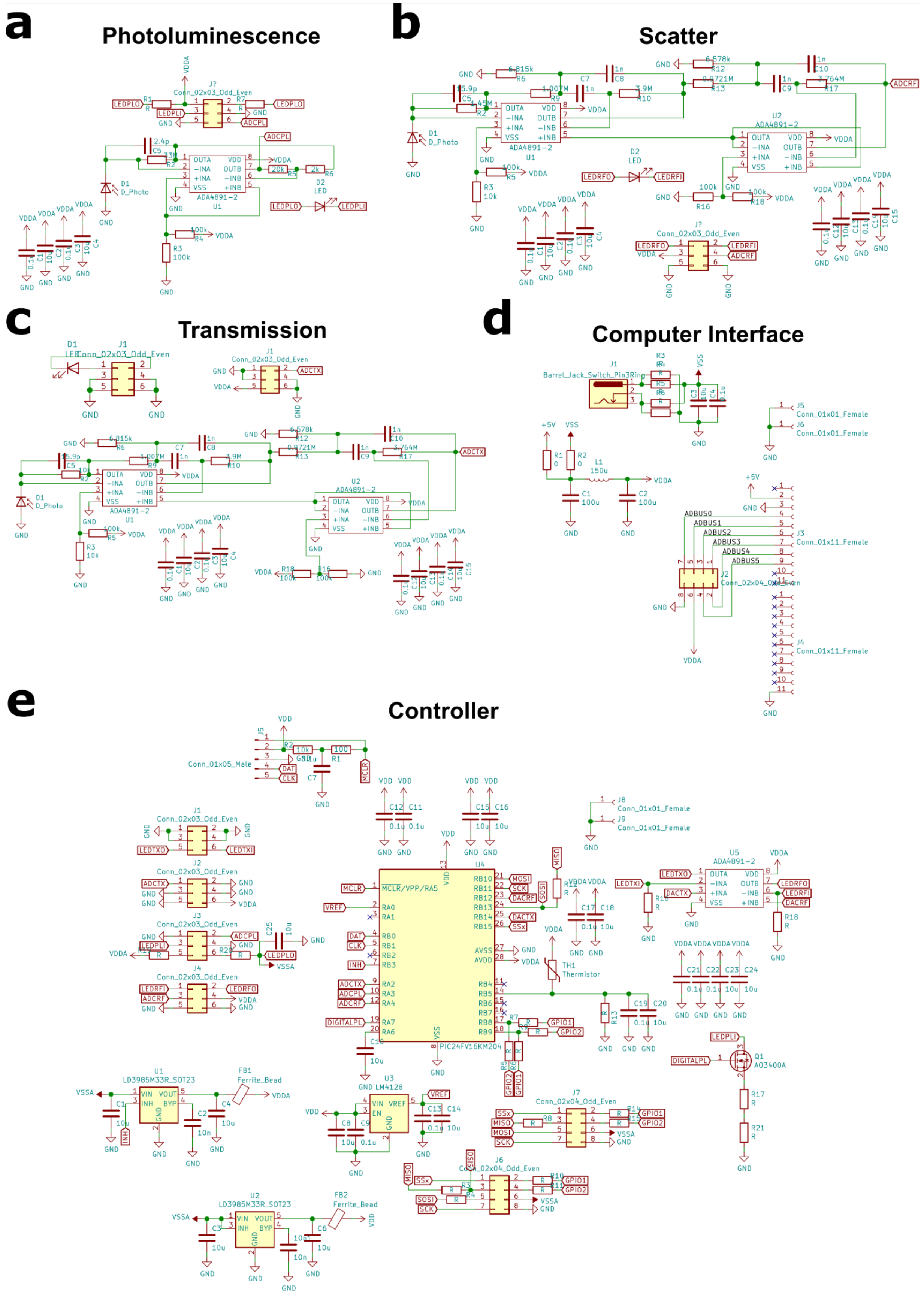


Supplementary Figure 2. Hardware schematics. (a) Photoluminescence hardware used in lifetime decay capture. (b) Scatter hardware to measure aggregate density. (c) Transmission hardware used in measuring phenol-red absorption-based pH. (d) Schematic design for computer interface to controller. (e) Control board schematic, encompassing the microcontroller and peripheral hardware.

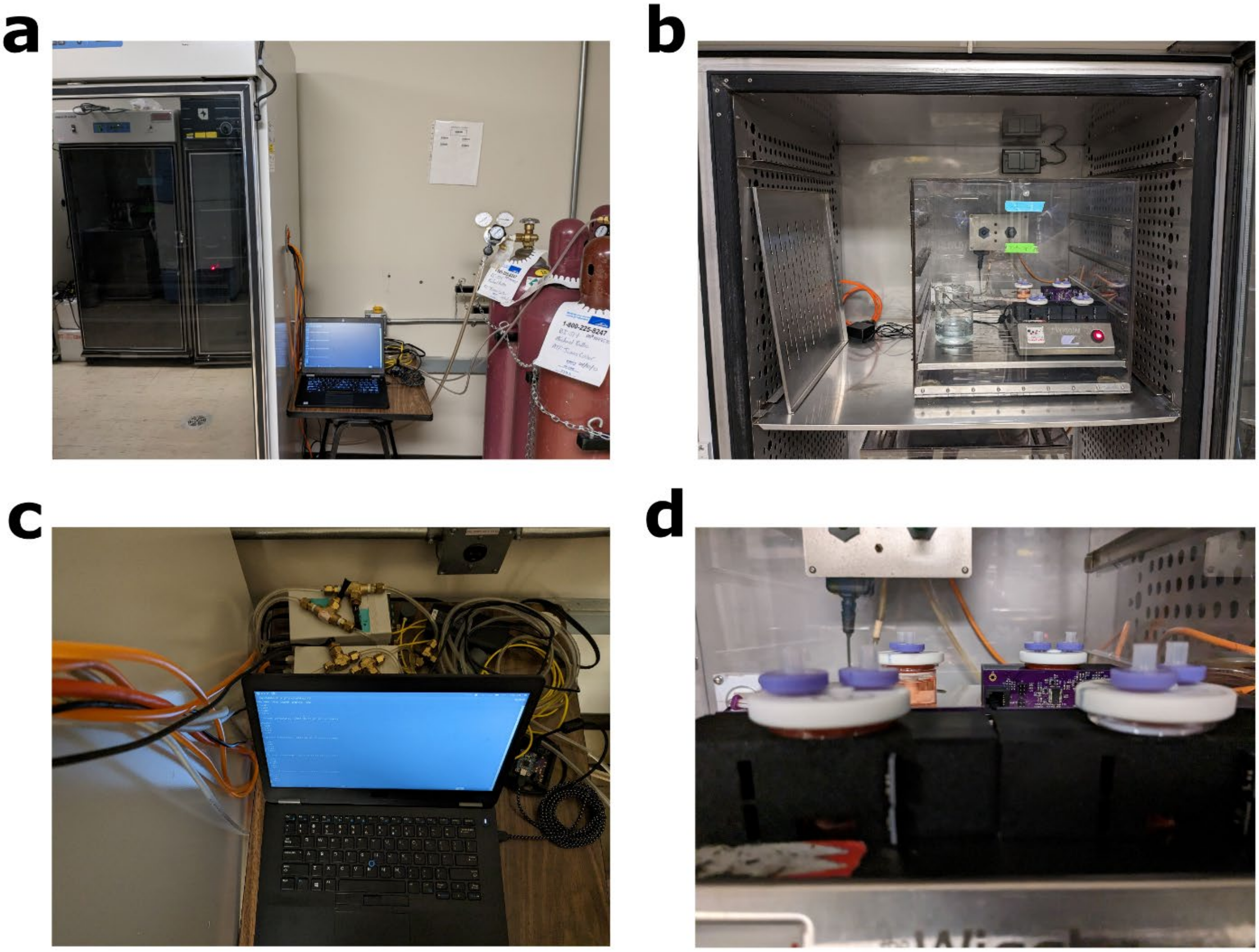


Supplementary Figure 3. Laboratory experimental setup. (a) Incubator positioning adjacent to compressed gas tanks, controllers and personal computer. (b) Incubator setup, with optoelectronics, mechanical frame, and scale-down reactors positioned inside of a hypoxia chamber. (c) Personal computer, interface board, and gas controllers for the hypoxia chamber. (d) Close-up of optoelectronic system inside the hypoxia chamber during operation.

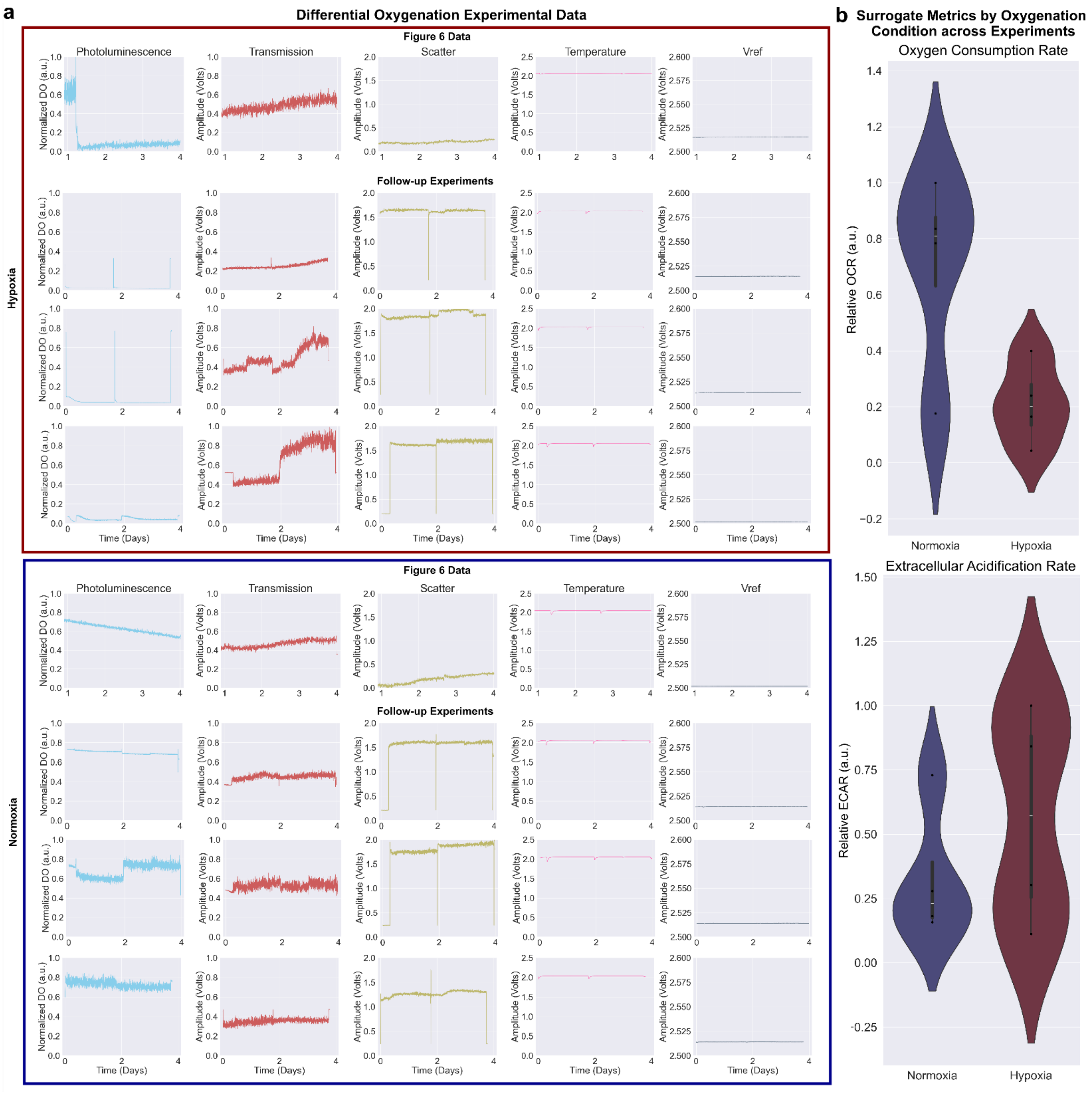


Supplementary Figure 4. (a) Experimental data obtained from initial study and follow-up differential oxygenation experiments. Follow-up experiments were carried out over 4 days, with offline sampling at days 2 and 4. (b) Relative OCR and ECAR were calculated and aggregated from these experiments.
